# A *Chryseobacterium massiliae* pore-forming MACPF domain protein mediates intra and interspecies competition against *Bacteroides*

**DOI:** 10.1101/2025.10.02.679979

**Authors:** Bianca Audrain, David Perez-Pascual, Stanislas Thiriet-Rupert, Joaquin Bernal-Bayard, Franziska Stressmann, Jean-Marc Ghigo

## Abstract

Microbiota play crucial roles in host health, including protection against pathogens through competitive interactions between commensal and pathogenic bacteria that are mediated by direct contact or secreted factors. We previously demonstrated that *Chryseobacterium massiliae*, a zebrafish commensal, protects larvae against infection by *Flavobacterium covae* (formerly *F. columnare*). Here, we investigated whether interference interactions contribute to this protective effect. We found that *C. massiliae* culture supernatant inhibits *F. covae* growth and a transposon mutagenesis screen identified mutants lacking this activity. All identified mutants carried insertions in a gene encoding a protein homologous to *Bacteroidales* BSAP pore-forming toxins, characterized by a Membrane Attack Complex/Perforin (MACPF) domain. We showed that this protein, which we named CSAP-1 (for *Chryseobacterium* Secreted Antimicrobial Protein) displays bactericidal, pore-forming activity that lyses *F. covae* cells. Unlike BSAP proteins from *Bacteroides* spp., CSAP-1 displays broader antagonistic activity, targeting multiple species across the *Flavobacterium* and *Chryseobacterium* genera - thus mediating interspecies and intergenus inhibition within *Bacteroidetes*. Although CSAP-1 is not essential for the *in vivo* protective effect of *C. massiliae*, administration of purified CSAP-1 alone confers significant protection to zebrafish larvae against sensitive *F. covae* infection.

This study therefore identifies CSAP-1 as the first MACPF protein from *C. massiliae* with broad-spectrum inhibitory activity against members of the order *Flavobacteriales*. These findings highlight CSAP-1 as a promising candidate for the development of novel antimicrobial strategies and warrant further mechanistic investigation.

## INTRODUCTION

Animal-associated microbiota are dense and dynamic communities that favorize interactions between microorganisms. Studies investigating the impact of bacteria-bacteria interactions on the composition and function of host-associated microbiota have highlighted the critical role of antagonistic interactions, often involving exploitative competition for shared resources or direct competitive interferences between bacteria (interference competition) (Boopathi et al. 2021; Sorbara and Pamer 2019; Wang et al. 2024). The use of controlled *in vivo* models led to significant progress in the characterization of the bacterial factors involved in competition against incoming pathogens, which were shown to contribute to microbiota-based infection resistance through niche exclusion and/or direct growth inhibition (Douglas 2019; Hecht et al. 2016; Leshem et al. 2020; Stagaman et al. 2020). Using a gnotobiotic zebrafish model of infection, we previously demonstrated that germ-free zebrafish larvae succumbed to the infection by the fish pathogen *Flavobacterium covae* (formerly *F. columnare*). By contrast, re-conventionalization with *Chryseobacterium massiliae* — a key member of the zebrafish microbiota — conferred complete protection against the pathogen (Stressmann et al. 2021). Moreover, *C. massiliae* was also shown to protect adult zebrafish as well as rainbow trout larvae from infection by *F. covae* (Perez-Pascual et al. 2021; Stressmann et al. 2021).

Considering that both bacteria belong to the *Bacteroidetes* phylum, the protection conferred by *C. massiliae* against *F. covae* could be due to competition for the same resources. Alternatively, *C. massiliae* could produce antimicrobial factors antagonizing *F. covae* growth or virulence. *Bacteroidetes* species such as *Bacteroides fragilis* were indeed previously shown to display antimicrobial activity affecting microbiota composition and pathogen colonization, either by cell-to-cell contact, via the type VI secretion system (T6SS) (Chatzidaki-Livanis et al. 2016; Coulthurst 2019; Coyne and Comstock 2019; Russell et al. 2014), or upon production of bacteriocins (Chatzidaki-Livanis et al. 2017; Coyne et al. 2019; Heilbronner et al. 2021; Jiang et al. 2024; Matano et al. 2021).

The characterization of competitive factors secreted by *Bacteroidales* also identified pore-forming proteins containing a Membrane Attack Complex/Perforin (MACPF) domain, first characterized in proteins of the eukaryote complement system and cytotoxic lymphocytes perforins (Brennan et al. 2010; Rosado et al. 2008). Four of such MACPF domain proteins called BSAP-1 to -4 for *<u>B</u>acteroidales*-<u>S</u>ecreted <u>A</u>ntimicrobial <u>P</u>rotein have been described (Chatzidaki-Livanis et al. 2014; McEneany et al. 2018; Roelofs et al. 2016; Shumaker et al. 2019) and *B. fragilis* BSAP-1 has been shown to alter membrane permeability via the formation of pores into the membrane of target bacteria (Chatzidaki-Livanis et al. 2014). These proteins are ubiquitous and responsible for intraspecies competition in *Bacteroides* species, providing an ecological advantage in each natural microbial community (Chatzidaki-Livanis et al. 2014; McEneany et al. 2018; Roelofs et al. 2016; Shumaker et al. 2019).

In this study, we aimed to identify the molecular factor involved in *C. massiliae* protection against *F. covae* infection in zebrafish. After developing random and targeted mutagenesis in *C. massiliae*, we identified a new MACPF domain-containing protein killing *F. covae* by cell lysis. By contrast with *Bacteroides* sp. BSAP proteins, we showed that this *C. massiliae* protein, renamed CSAP-1 by homology with BSAP proteins, mediates both interspecies and intergenus inhibitory activity on several *Flavobacterium* and *Chryseobacterium* species. Finally, whereas CSAP-1 is not required for the protection conferred by *C. massiliae* against *F. covae* infection *in vivo*, we determined that adding purified CSAP-1 protein protects axenic zebrafish larvae against *F. covae* infection. These findings position CSAP-1 as a promising candidate for alternative early-stage antimicrobial strategies to treat *F. covae* infection.

## RESULTS

### *Chryseobacterium massiliae* secretes a MACPF domain protein inhibiting the growth of the fish pathogen *F. covae*

To investigate the mechanisms underlying *C. massiliae* protection against *F. covae* infection, we tested the ability of *C. massiliae* to inhibit *F. covae* ALG-00-530 (*F. covae*^ALG^). We showed that *C. massiliae* bacteria-free overnight supernatant spotted on agar plate overlayed with *F. covae*^ALG^ displayed an inhibitory activity (Fig. 1A). To identify the inhibitory molecule(s) secreted by *C. massiliae*, we performed a random transposon mutagenesis using the pHimarEm1 plasmid, delivering *Himar1* transposons (Braun et al. 2005), and screened the activity of the supernatant of ca. 4000 *C. massiliae* transposon mutants using agar overlay spot assay (Supplementary Fig. S1). This enabled us to identify four mutants that had lost the ability to inhibit *F. covae*^ALG^ growth (Fig. 1B). All four transposons were inserted into the *bc7_v1_410027* gene encoding a protein with a membrane attack complex/perforin (MACPF) domain (Fig. 1C). To confirm that the transposon insertion into *C. massiliae bc7_v1_410027* was responsible for the loss of growth inhibition against *F. covae*^ALG^, we used genetic tools developed for other *Bacteroidetes* species (Zhu et al. 2017) to generate a full *bc7_v1_410027* mutant in *C. massiliae* and we confirmed that the resulting mutant strain *C. massiliae Δbc7_v1_410027* lost its ability to inhibit *F. covae*^ALG^ growth (Fig. 1B).

**Figure 1:**
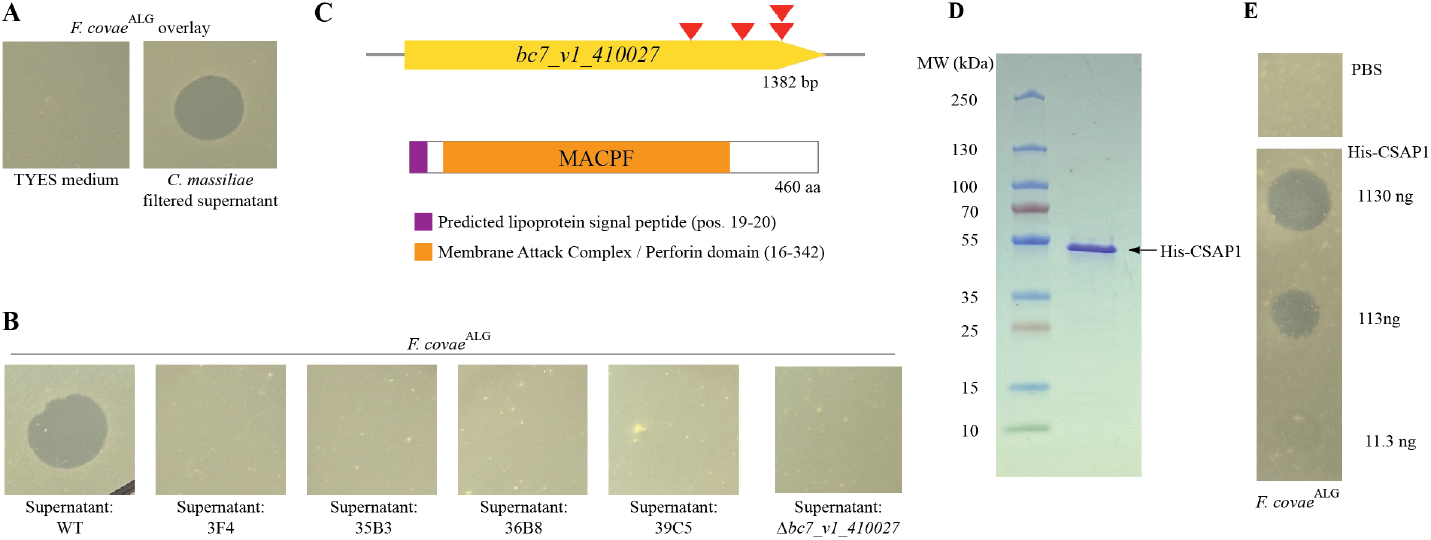
*C. massiliae* secretes a MACPF domain containing protein able to inhibit the growth of *F. covae*^ALG^. (**A**) Overlay assay showing that *C. massiliae* secretes a factor that inhibit the growth of *F. covae*^ALG^. 10 µl of TYES media or filtered supernatant of a *C. massiliae* culture were dropped on *F. covae*^ALG^ lawn. (**B**) Supernatant of four transposon mutants and *bc7-v1-410027* deletion mutant in *C. massiliae* lost their ability to inhibit the growth of *F. covae*^ALG^. (**C**) Localization of transposon insertion into *bc7_v1_410027*. Arrows indicate insertion of the four transposons. MACPF domain was identified on InterPro while signal peptide was identified using Signal6.0. (**D**) Coomassie stained gel of purified His-CSAP-1. Lane 1: PageRuler Prestained plus ladder (Thermo #26619); Lane 2: Purified His-CSAP-1 after dialysis (see Material and Methods). (**E**) Agar overlay assay showing inhibitory activity displayed by purified His-CSAP-1 on overlaid *F. covae*^ALG^ whereas PBS has no impact. Amounts of protein in 10 µL are indicated.

Whereas all our attempts to complement the *Δbc7_v1_410027* mutant strain failed, we showed that a purified His-tagged version of the bc7_v1_410027 protein inhibited *F. covae*^ALG^ growth (Fig. 1DE). These results demonstrated that the *C. massiliae* antimicrobial activity against *F. covae*^ALG^ does not require direct cell contact and is mediated by the secretion of the protein encoded by *bc7_v1_410027*.

While this bc7_v1_410027 protein has little sequence similarity with BSAPs identified from *Bacteroides spp*., it displays critical MACPF residues as well as a predicted SpII cleavage site in their signal sequence (Chatzidaki-Livanis et al. 2014) (Supplementary Fig. S2). We therefore proposed to rename this protein CSAP-1, for *Chryseobacterium* Secreted Antimicrobial Protein, and its corresponding gene *csap-1*.

### *C. massiliae* CSAP-1 mediates interspecies antagonism over other *Bacteroidetes*

To explore the host range of *C. massiliae* CSAP-1 activity in zebrafish, we used the agar overlay spot assay to test the activity of purified His-CsaP-1 and WT *C. massiliae* filtered supernatant on the nine core Proteobacteria identified in zebrafish larvae microbiota (Stressmann et al. 2021). Our analysis showed that CSAP-1 did not affect the growth of any of the tested strains (Supplementary Fig. S3). We then expanded our analysis to different *Bacteroidetes* bacteria including *Flavobacterium* spp., *Bacteroides* spp. and *Chryseobacterium* spp.. We showed that the growth of 3 out 9 tested *F. covae* and 15 out 22 tested *F. columnare* strains, all originally classified as *F. columnare* the causative agent of columnaris (LaFrentz et al. 2022), were inhibited by CSAP-1 without any specificity to host and geographic origins nor zebrafish virulence (Table 1). We also identified that *Flavobacterium psychrophilum* and *Flavobacterium branchiophilum* were inhibited by CSAP-1 whereas *Flavobacterium davisii* (2), *Flavobacterium ceti* (1), *Elizabethkingia meningoseptica* as well as two *Bacteroides* species (*B. thetaiotaomicron* and *B. ovatus*) did not reveal any sensitivity to CSAP-1. We then used the *C. massiliae* supernatant and the purified His-CSAP-1 protein to test the CSAP-1 intra-genus inhibitory activity and showed that CSAP-1 was active against *C. defluvii, C. indoltheticum, C. balustinum, C. scopthalmum* and several *Chryseobacterium sp*. (Table 1). It should be noted that the supernatant of *C. massiliae Δcsap1* also inhibited the growth of different *Chryseobacterium* species including *C. defluvii, C. gleum, C. massiliense, C. piscium and some Chryseobacterium sp*., suggesting that *C. massiliae* produces diverse antimicrobial molecules (Table 1).

**Table 1:**
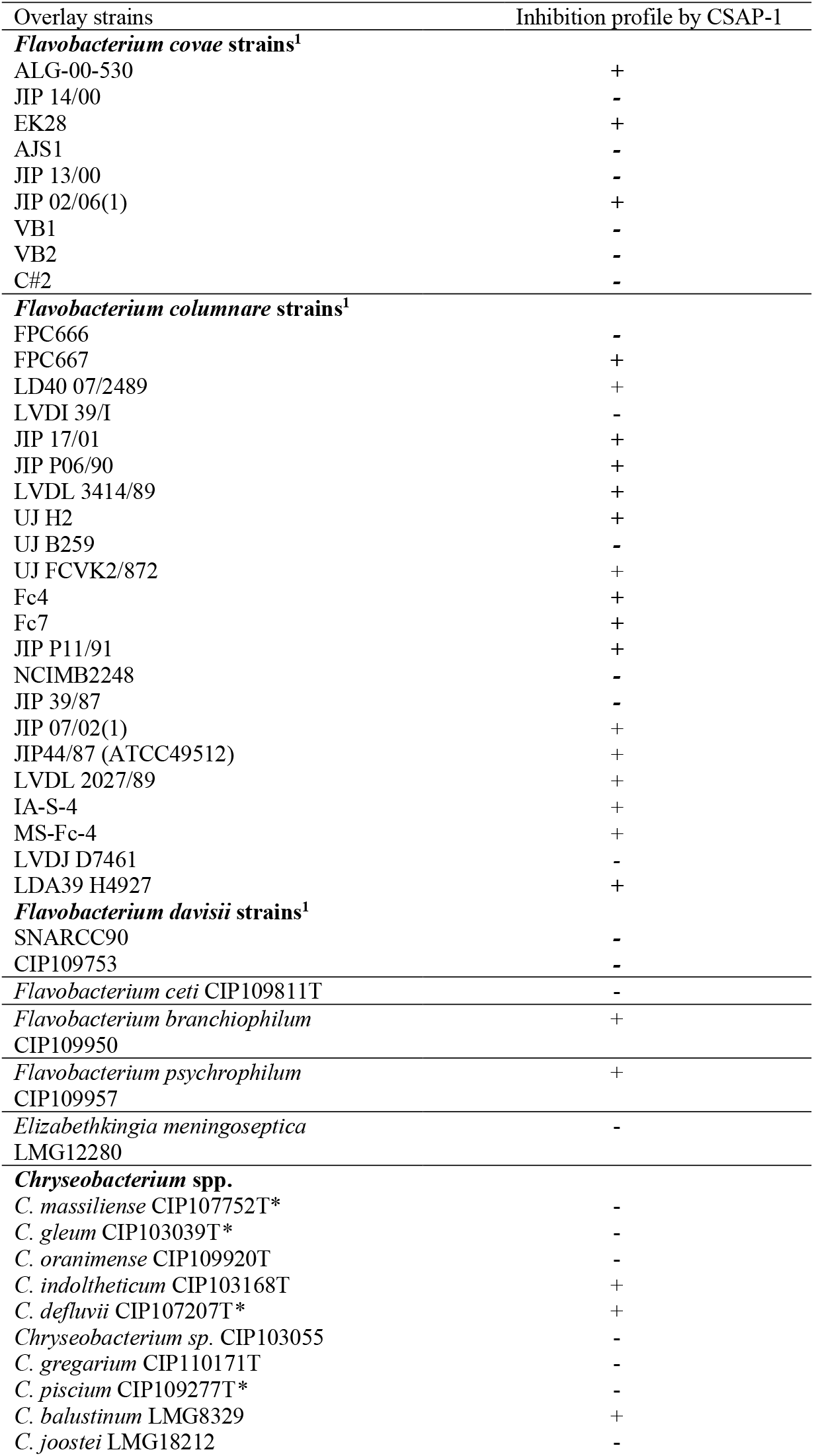
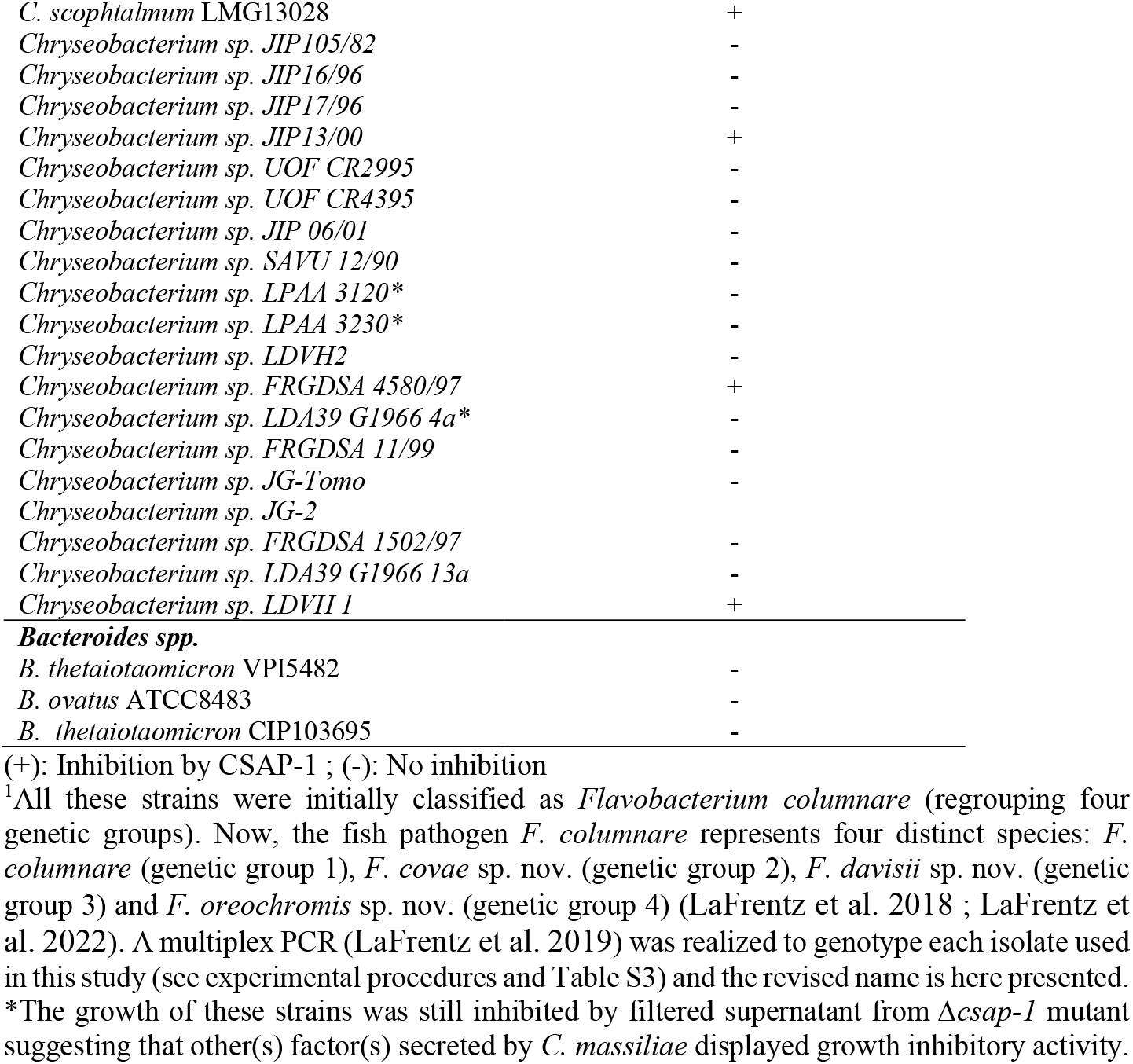
Analysis of growth inhibition (agar overlay test) mediated by CSAP-1 among *Bacteroidetes*.

Taken together, our data indicated that, by contrast with the intraspecies antimicrobial activity of *Bacteroides* BSAP MACPF proteins (Chatzidaki-Livanis et al. 2014; McEneany et al. 2018; Roelofs et al. 2016; Shumaker et al. 2019), *C. massiliae* CSAP-1 mediates both intra- and extra-genus antagonistic interactions.

### CSAP-1 kills *F. covae*^ALG^ by cell lysis

To investigate the mechanism by which CSAP-1 inhibits the growth of *F. covae*^ALG^, we quantified *F. covae*^ALG^ survival after 30 min of exposure to purified His-CSAP-1 protein as well as to filtered supernatant either from WT *C. massiliae* or its corresponding *Δcsap-1* mutant. We observed a reduction in *F. covae*^ALG^ viability after exposure to His-CSAP-1 or WT supernatant, whereas the survival was unaffected by the exposure to *C. massiliae Δcsap-1* supernatant, suggesting that CSAP-1 displayed a bactericidal activity in these conditions (Fig. 2A). We then investigated the potential pore-forming activity of CSAP-1 by monitoring the uptake of propidium iodide into *F. covae*^ALG^ exposed to CSAP-1. This red-fluorescent dye can only cross compromised bacterial membranes and we observed *F. covae*^ALG^ cell lysis after 30 min exposure to *C. massiliae* wild-type supernatant as demonstrated by an increased labeling of red fluorescence in bacteria and the presence of burst cell and envelope debris (Fig. 2B). By contrast, no modifications of fluorescence levels or cell morphology were observed when bacteria were exposed to *Δcsap-1* supernatant (Fig. 2B).

**Figure 2:**
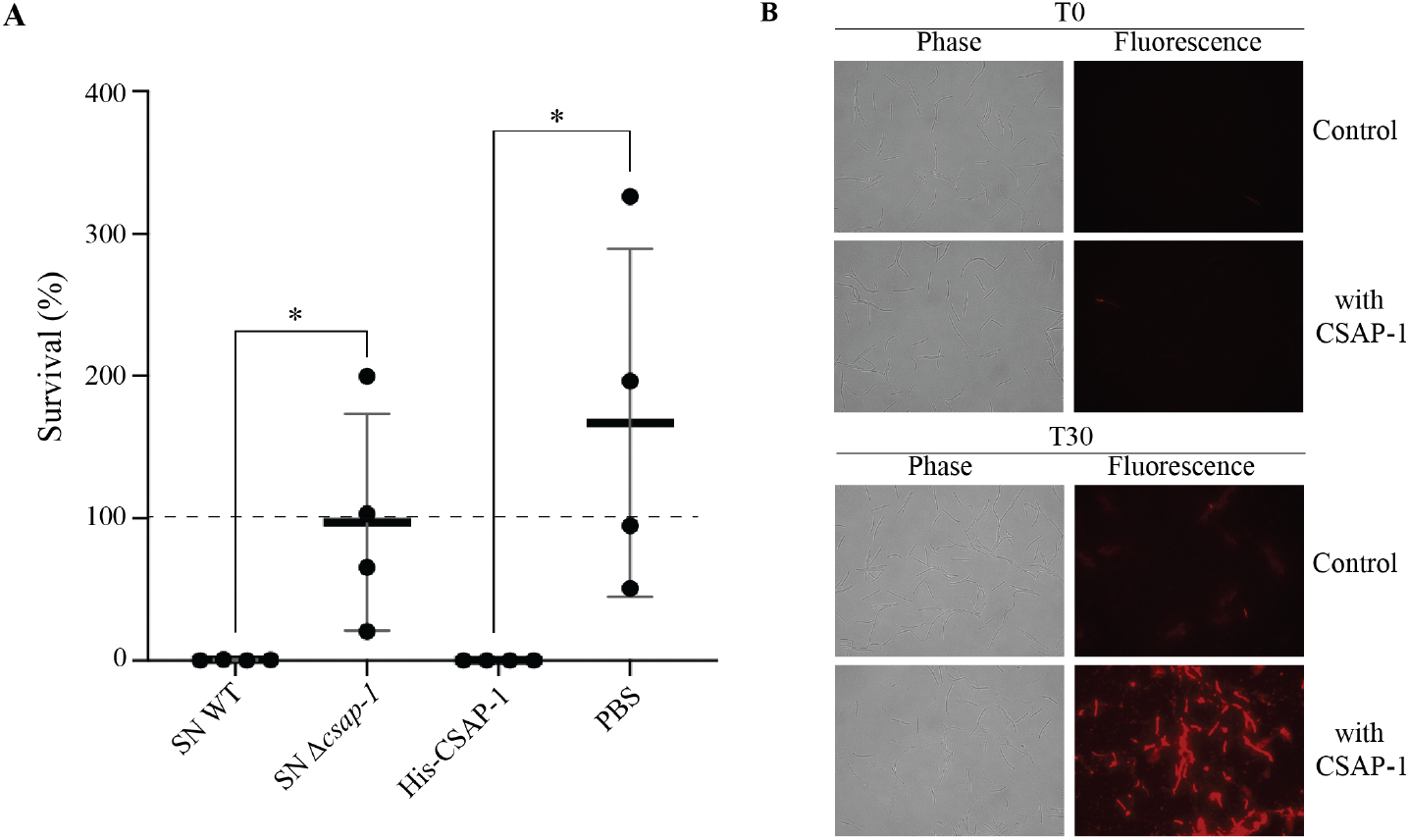
CSAP-1 can lyse *F. covae*^ALG^ cells. **(A)***F. covae*^ALG^ cells were exposed either to filtered supernatant of WT or of *Δcsap-1* mutant strain (*left*), or to purified His-CSAP-1 or PBS (*right*). Quantification of survival was determined by CFU counting upon 30 min of exposure and expressed as percent of survival compared to CFU at T0. Statistics correspond to unpaired, non-parametric Mann-Whitney test. *p<0.05. (**B**) Microscopic analysis of propidium iodide incorporation by *F. covae*^ALG^ cells exposed to filtered supernatant of WT *C. massiliae* (CSAP-1) or *Δcsap-1* (Control). Images are shown at the starting point (before exposure) and 30 min after exposure to each supernatant.

### Purified CSAP-1 protects zebrafish larvae against *F. covae* infection

To determine whether the CSAP-1-dependent ability of *C. massiliae* to inhibit and kill *F. covae*^ALG^ is involved in zebrafish protection against *F. covae* infection (Stressmann et al. 2021), we reconventionalized germ-free zebrafish larvae at 4 days post-fertilization (dpf) with either WT *C. massiliae* or its corresponding *Δcsap-1* mutant prior to exposing them to *F. covae*^ALG^. We showed that both strains provide protection to otherwise lethal doses of the pathogen, indicating that CSAP-1 secretion does not contribute to *C. massiliae* zebrafish protection against *F. covae* infection (Fig. 3).

**Figure 3:**
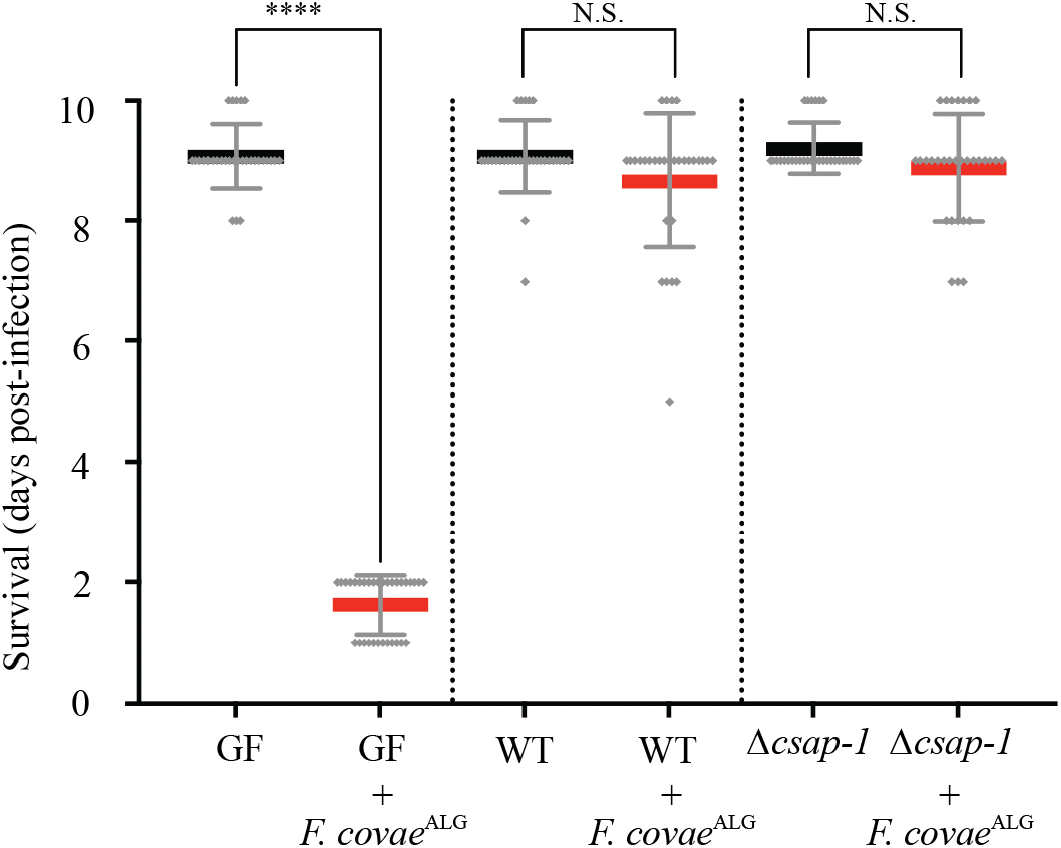
CSAP-1 does not contribute to *C. massiliae* zebrafish protection against *F. covae*^ALG^ infection. Zebrafish larvae were reconventionalized at 4 dpf with either *C. massiliae* WT strain or its corresponding *csap-1* deletion mutant and were infected at 6 dpf with *F. covae*^ALG^ (red). Larvae mortality was monitored daily, and surviving fish were euthanized at day 9 post-infection. Mean survival is represented by a thick horizontal bar with standard deviation. Each point represents a single fish, and each condition was repeated at 3 independent times (here, n = 30-32 fish per condition). GF = Germ-Free. Black mean corresponds to non-infected larvae. Statistics correspond to unpaired, non-parametric Mann-Whitney test. **** p<0.0001; N.S.: non-significant.

However, exposing infected larvae to purified CSAP-1 could nevertheless protect them against *F. covae* infection. To test this, we transferred zebrafish larvae after 3 hours of immersion with the pathogen, in water containing either PBS or purified His-CSAP-1 (8 µg) and we monitored the survival for 9 days. As a negative control, we also infected zebrafish larvae with the virulent but CSAP-1 – resistant *F. covae* strain C#2. We observed that exposure to purified CSAP-1 allows larvae to survive infection by sensitive *F. covae*^ALG^ but not by the resistant one (Fig. 4A). We determined that whereas 366 +/-70.7 colony forming units (cfu) were recovered from non-treated infected larvae, almost no *F. covae*^ALG^ cells were detected when exposed to His-CSAP-1 after infection (1 cfu) (Fig. 4B). These data therefore showed that, whereas the production of CSAP-1 is not a key determinant of *C. massiliae* protection against *F. covae*, exposure to CSAP-1 alone can prevent infection against CSAP-1-sensitive strains.

**Figure 4:**
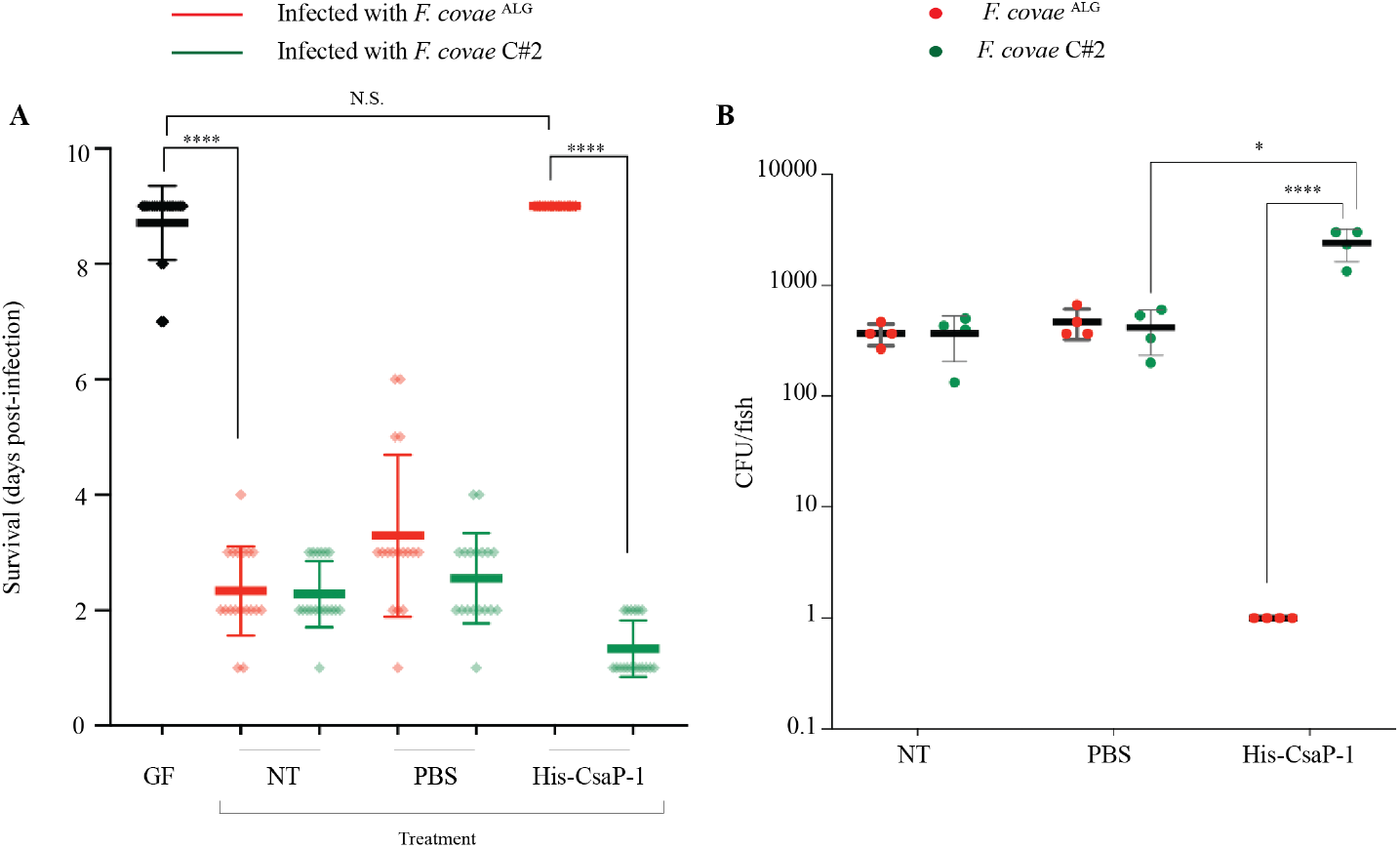
Exposure to His-CSAP-1 allows zebrafish larvae to survive infection by CSAP-1-sensitive strain of *F. covae*. (**A**) 6 dpf germ-free zebrafish larvae were exposed 3h to CSAP-1 sensitive (red) or resistant (green) strain of *F. covae* using bath immersion before transfer into sterile water (NT, non-treated), or sterile water containing either PBS or His-CSAP-1 (8 µg). Larvae mortality was monitored daily and surviving fish were euthanized at day 9 post-infection. Mean survival is represented by a thick horizontal bar with standard deviation. Each point represents a single fish (n = 16-21 fish per condition) and each condition was repeated at 2 independent times. GF = Germ-Free. Black mean corresponds to non-infected larvae. Statistics correspond to unpaired, nonparametric Mann-Whitney test. ****p<0.0001; N.S. or absence of *: non-significant. (**B**) 6 dpf germ-free zebrafish larvae were exposed 3h to CSAP-1 sensitive (red) or resistant (green) strain of *F. covae* using bath immersion before transfer into sterile water (NT, non-treated), or sterile water containing either PBS or His-CSAP-1. GF zebrafish colonization of *F. covae* sensitive (ALG) or resistant (C#2) to CSAP-1 was measured by CFU per fish 24 hours after transfer into the different conditions. Each point represents a single fish where fill color indicates the strain that was counted.

## DISCUSSION

In multispecies bacterial communities, individual species may coexist harmoniously or engage in competition for resources, either indirectly or antagonizing other bacteria via cell-contact-dependent mechanisms (Aoki et al. 2010), or using strategies involving the release of antibacterial compounds (Peterson et al. 2020). Here, we showed that CSAP-1, a MACPF domain-containing protein secreted by *C. massiliae*, displayed interspecies and intergenus inhibitory activity. CSAP-1 affects several species belonging to the *Flavobacteriales* order, including pathogenic *F. covae* and *F. columnare* (originally regrouped as *F. columnare*), the causative agents of columnaris disease in wild and cultured fish populations worldwide (Declercq et al. 2013).

Although BSAPs and CSAP-1 share little homology, they display key conserved MACPF residues shown to be necessary for inhibitory activity in BSAP-1, a pore-forming protein permeabilizing membranes (Chatzidaki-Livanis et al. 2014), a physiology-disrupting event leading to bacterial cell death. Consistently, we showed that CSAP-1 from *C. massiliae* induces rapid propidium iodide labeling and *F. covae*^ALG^ cell lysis, indicative of a pore-forming activity. BSAPs and CSAP-1 also both have a predicted SpII cleavage site in their signal sequence suggesting they all may be released through the production of outer membrane vesicles (Chatzidaki-Livanis et al. 2014). BSAPs proteins were shown to target specific surface molecules such as ß-barrel outer membrane protein (BSAP-1 and BSAP-4), or lipopolysaccharide glycans (BSAP-2 and BSAP-3) (Chatzidaki-Livanis et al. 2014; McEneany et al. 2018; Roelofs et al. 2016; Shumaker et al. 2019). Interestingly, BSAP encoding genes have been shown to be adjacent to genes producing a functional equivalent of their target molecules in the sensitive strains. This specific localization allowed bioinformatic analyses to identify resistance or sensitivity determinants. We showed here that CSAP-1 affects the growth of 15 out 22 tested *F. columnare* strains and 3 out 9 tested *F. covae*, all initially classified as *F. columnare* strains, without any specificity to geographic or historical host origins. We attempted to identify CSAP-1 target by comparing the genomic sequences of 7 CSAP-1 sensitive strains *versus* 7 resistant ones, but we could not identify any gene(s) present only in the sensitive strains and absent in the resistant strains (see details in Experimental procedures and Supplementary data 1). Further research will therefore be required to identify potential targets of CSAP-1.

Interestingly, the action spectrum of CSAP-1 is not restricted to the causative agent of columnaris (*F. covae, F. columnare*) since other *Flavobacterium* species are sensitive to CSAP-1, such as the fish pathogens *F. psychrophilum* and *F. branchiophilum*, as well as some species belonging to the genus *Chryseobacterium* such as *C. defluvii, C. indoltheticum* and several *Chryseobacterium* sp. isolated from the aquatic environment. Whereas we did not detect any inhibitory activity against the zebrafish larvae core microbiota, mostly belonging to the *Proteobacteria* phylum (Stressmann et al. 2021), the production of CSAP-1 may confer a competitive advantage at a later developmental stage, since the adult zebrafish microbiota includes diverse taxonomic groups within *Bacteroidetes* (Roeselers et al. 2011; Stephens et al. 2016).

*C. massiliae* was shown to protect GF zebrafish larvae against *F. covae* infection (Stressmann et al. 2021) but our results showed that CSAP-1 is not necessary for this protection since *C. massiliae Δcsap-1* mutant is still protective. This suggests that the protection driven by *C. massiliae* against *F. covae* infection is not exclusively associated with CSAP-1 production. However, GF zebrafish larvae exposed to purified CSAP-1 survived *F. covae*^ALG^ infection, suggesting that CSAP-1 could still contribute to protection. Currently, the prevention of columnaris disease relies heavily upon using good aquaculture management practices to reduce risk factors, vaccination, or the excessive use of antibiotics (Declercq et al. 2013) (Pulkkinen et al. 2010). In this context, CSAP-1 could be considered a promising compound to treat the early stages of infection by *F. covae*. Future studies should explore the efficacity and dynamic of CSAP-1 to treat *F. covae* or *F. psychrophilum* infections in more epidemiologically relevant fish species.

In conclusion, this study reports the first identification and characterization of *C. massiliae* CSAP-1, a MACPF-containing protein exhibiting interspecies inhibitory activity against several members of the order *Flavobacteriales* and significantly enhancing zebrafish survival during early-stage of *F. covae* infection. While further studies will clarify its mechanism of action, these findings highlight CSAP-1 as a promising candidate for alternative antimicrobial strategies.

## EXPERIMENTAL PROCEDURES

### Bacterial strains, plasmids and growth conditions

Bacterial strains and plasmids used in this study are listed in Table S1. *Flavobacterium* spp. and *Chryseobacterium* spp. strains were grown at 28°C in tryptone yeast extract salts (TYES) broth [0.4 % (w/v) tryptone, 0.04 % yeast extract, 0.05 % (w/v) MgSO_4_ 7H_2_O, 0.02 % (w/v), CaCl_2_ 2H_2_O, 0.05 % (w/v) D-glucose, pH 7.2] whereas *E. coli* strains were grown in Luria-Bertani (LB) medium at 37°C. If required, 15 g/L of agar was added for solid medium. When necessary, antibiotics were added at the following concentration: Erythromycin (100 µg/mL), Ampicillin (100 µg/mL).

### Multiplex PCR for genotyping *F. columnare* and revision of species designation

In recent years, LaFrentz *et al*. identified that the fish pathogen *F. columnare* represents four distinct species depending on its genotype (LaFrentz et al. 2018; LaFrentz et al. 2022). Genotyping of the 33 isolates used in our study was performed using their multiplex PCR (LaFrentz et al. 2019). First, genomic DNA was extracted from overnight cultures using the Wizard Genomic DNA Purification Kit (Promega) following the instructions for Gram-negative bacteria and 20 ng were then used as PCR template. The PCR was performed with Phusion Flash High-Fidelity PCR Master Mix (Thermofisher Scientific, F548) and the primer concentrations in the 25 μL reactions were 0.5 μM GG-fwd, 0.1 μM GG1-rev, 0.45 μM GG2-rev, 0.45 μM GG3-rev and 0.3 μM GG4-rev (Table S2). PCR amplification was realized using the following cycling protocol: one cycle at 95°C for 5 min; 40 cycles of 30 s at 94°C, 20 s at 56°C and 60 s at 72°C and a final cycle of 10 min at 72°C. PCR products were resolved on a 2% (w/v) agarose gel and sizes corresponding to each species were 415, 320, 894 and 659 bp for respectively *F. columnare* (genetic group 1), *F. covae* (genetic group 2), *F. davisii* (genetic group 3) and *F. oreochromis* (genetic group 4) (Fig. S4). Genetic groups determined by this multiplex PCR and respective reclassification are presented in Table S3.

### Agar overlay assay for growth inhibition detection

The growth inhibitory effect of cell-free supernatant has been evaluated using an agar spot test. Briefly, 125 μL from an overnight culture of targeted cells adjusted to OD 1 were mixed to 5 mL of top agar (0.6-0.7% agar) and overlaid on plates. Overnight culture of *C. massiliae* was centrifuged for 5 min at 8000 rpm and resulting supernatants were filtered through 0.22 µm filters. 5 µL of cell-free spent supernatant were then spotted on the overlay of targeted bacteria. Recipient strains were grown in appropriate liquid and corresponding top agar medium.

### Genetic manipulation in *C. massiliae* and *F. columnare*

#### (i) Transposon mutagenesis in *C. massiliae*

pHimarEm1, first described as a tool for random transposon mutagenesis in *Flavobacterium johnsoniae* was introduced into *C. massiliae* by conjugation from *E. coli* S17λpir (Braun et al. 2005). Briefly, 1 mL of the donor strain grown in LB was washed twice in TYES whereas 2 mL from *C. massiliae* culture grown in TYES were washed twice in TM buffer (20 mM Tris-HCl, 20 mM MgSO_4_, pH 7.2). Both were resuspended in 50 µL of TYES, pooled together with a 1:2 donor-recipient ratio and spotted on a 0.45 µm filter on TYES plate. After 24 hours of incubation at 28°C, spots were resuspended in TYES and transconjugants were selected on TYES plates containing 100 µg/mL erythromycin. Isolated colonies were resuspended in 100 µL of TYES supplemented with erythromycin in 96-well plates and grown 24 hours to establish the library regrouping 4000 mutants.

#### (ii) Creation of deletion mutants in *C. massiliae*

pYT313, a suicide vector conferring erythromycin resistance and containing *sacB* gene controlled by *ompA* promoter from *F. johnsoniae* for counter selection (Zhu et al. 2017) was used to create deletion mutants in *C. massiliae*. 1 kb region upstream and downstream of the gene of interest, and vector were amplified by PCR using Phusion Flash High-Fidelity PCR Master Mix (Thermofisher Scientific, F548) and primers listed in Table S2. Ligation of the three fragments was then performed by Gibson assembly: inserts and vector were mixed with Gibson master mix 2X (100 μl 5X ISO buffer, 0.2 μl 10,000 U/ml T5 exonuclease [NEB catalog number M0363S], 6.25 μl 2,000 U/ml Phusion HF polymerase [NEB catalog number M0530S], 50 μl 40,000 U/ml *Taq* DNA ligase [NEB catalog number M0208S], and 87 μl dH_2_O for 24 reactions) and incubated at 50°C for 30 min. The resulting mix was first introduced into *E. coli* S17λpir, a donor strain then used to transfer the vector to recipient strains by conjugation (see above). After 48 hours of incubation at 28°C, spots were resuspended in TYES, serially diluted and plated on TYES supplemented with 200 µg/mL erythromycin to select clones that had undergone the first event of recombination (plasmid insertion into the chromosome). After 2 days at 28°C, clones were streaked once on TYES plus erythromycin and then grown overnight in TYES without antibiotics to allow the second event of recombination. 100 µL of overnight culture were plated on TYES plates containing 5% sucrose to select for loss of the plasmid. According to the first insertion site, the second event of recombination led to either the deletion of the targeted gene or a wild-type version. Clones were streaked once on TYES with sucrose before being isolated on TYES plates. Gene deletion was confirmed by PCR and sequencing of the region (see primers in Table S2).

### Construction and purification of His-tagged CSAP-1

An N-terminal His-tagged fusion of CSAP-1 was constructed and produced as following. *csap-1* (corresponding to 21-461 amino acids) was cloned into pET16b by Gibson assembly (see above) using primers listed in Table S2. The resulting protein, His-CSAP-1, was induced from a culture in exponential phase by adding 0.1 mM IPTG for 3 h at 30°C. His-CSAP-1 was purified using HisLink Spin Protein Purification System kit (Promega) and eluted protein was dialyzed in PBS overnight. Protein concentration was determined using the Qubit Fluorometer (Q33211, with Qubit Protein assay kit, Invitrogen). Protein expression and purification were also checked using western blot detection and/or visualization on blue Coomassie stained gel. Different quantities of purified His-CSAP-1 were spotted on the overlay of targeted bacteria prepared as previously mentioned (see above).

### Blue Coomassie staining

An 0.2 OD_600_ equivalent quantity of cells or purified protein in 1X Laemmli buffer (#1610747, Biorad) were boiled at 100°C for 5 min before being separated by SDS-polyacrylamide denaturing gel electrophoresis using precast TGX 4-15 % gradient gels (Bio-Rad). Gels were fixed one hour in 50% ethanol – 10% acetic acid, then stained with Blue Coomassie for one hour and finally incubated in 10% acetic acid for 4 x 30 min.

### *in vivo* experiments

Sterilization of zebrafish eggs as well as procedures to handle and raise germ-free and reconventionalized zebrafish larvae including feeding with *Tetrahymena thermophila* protozoans were performed as previously described (Stressmann et al. 2021). *(i) Reconventionalization*. After hatching, at 4 days post-fertilization (dpf), zebrafish larvae were reconventionalized by immersion with wild-type or *csap-1* mutant strains of *C. massiliae* as following. From overnight cultures in TYES broth at 28°C, bacteria were pelleted and washed twice in sterile water, then adjusted to OD_600_ = 1. Culture flasks containing germ-free fish were inoculated with bacteria at a final concentration of 5.10^5^ cfu/mL. *(ii) Infection with F. covae*. Reconventionalized zebrafish larvae were infected at 6 dpf for 3h by immersion in culture flasks with *F. covae* at a final concentration of 5.10^2^ cfu/mL. After infection, larvae were transferred into individual wells of 24-well plates and mortality was monitored daily. *in vivo* experiments were stopped at day 9 post-infection using tricaine (MS-222). Each condition regrouped 8 to 12 zebrafish larvae and independent experiments was repeated 2 to 3 times.

### Determination of bactericidal activity

From overnight cultures, *F. covae* was adjusted to OD 0.5 in fresh TYES medium and then co-incubated in a 1:1 ratio with either spent supernatant of wild-type *C massiliae* or the corresponding *csap-1* mutant. The same experiment was also performed using purified His-tagged CSAP-1. To determine *F. covae* survival after exposure to CSAP-1, colony-forming units of *F. covae* were monitored by plate counting on TYES agar.

### Propidium iodide labelling

*F. covae*^ALG^ adjusted to OD 0.5 in fresh medium was co-incubated with an equal volume of filtered supernatant of overnight cultures of WT *C. massiliae* or *csap-1* mutant. After 30 min, propidium iodide was added at 10 µg/mL and 2 µL of the mix were placed on a 0.8% agarose pad. Bacteria were immediately imaged by epifluorescence microscopy. *F. covae*^ALG^ adjusted to OD 0.5 in fresh medium before incubation with supernatant was also imaged as T0 control.

### Whole genome sequencing

Genomic DNA was extracted from overnight cultures using the Wizard Genomic DNA Purification Kit (Promega). Illumina whole-genome sequencing was performed by the Plateforme de Microbiologie Mutualisée (P2M) of Institut Pasteur and the genomes were assembled using SPAdes v3.13.0 (PMID 22506599) and annotated using RASTtk (25666585, 27899627) on the patricbrc.org database (31667520).

### *in silico* search for potential CSAP-1 targets

To identify potential CSAP-1 targets, we combined four *in silico* strategies based on genome comparisons: (i) we compared all genes from two sensitive and four resistant *Chryseobacterium* strains as well as seven sensitive and seven resistant *Flavobacterium* strains (see Supplementary Data 1 for strain names and related genome assembly accession numbers). (ii) As a more targeted approach, we also included to the genome comparisons the gene encoding the target of BSAP-1 and BSAP-2 from *B. fragilis* and *B. uniformis*, respectively (25339613, 27555309). (iii) Since the target of previously described MACPF-domain containing proteins can be encoded by neighbouring genes to the toxin-encoding gene, 10 genes upstream and 10 genes downstream of the CSAP-1 encoding gene were retrieved from *C. massiliae* genome and added to the analysis. (iv) Finally, because the LPS is known to be targeted by such toxins (30065309), the complete O-PS locus from *F. psychrophilum* (31139169) was also added to the analysis.

All genomes and gene sets were used as input to compute every possible group of orthologous genes (orthogroups) with OrthoFinder v2.3.8 (31727128). We then looked for patterns where genes belonging to a given orthogroup were present in all sensitive strains and absent from all resistant strains or the opposite. Such pattern would highlight genes either encoding for a potential target or involved in the target modification. Unfortunately, no specific pattern was identifiable by comparing sensitive and resistant strains (Supplementary Data 1).

## Data availability

All genomes sequenced and related annotation used in this study were deposited in NCBI under the Bioproject accession number PRJNA1302412. The assembly accession number for each genome can be found in Supplementary Data 1.

Input and output data for the orthogroup analysis are available at https://github.com/Sthiriet-rupert/CSAP-1.

## Statistical analyses

Statistical analysis was performed using Prism10 (GraphPad Software, Inc). We used only the non-parametric Mann-Whitney test. A cutoff P value of 5% was used for all tests (*, p<0.05; ****, p<0.0001).

## ACKNOWLEDGEMENTS

We are grateful to Mark McBride and Lionel Cladière for providing pHimar vector used in this study to perform random mutagenesis. We are very thankful to Mark McBride for providing several *Flavobacterium* sp. strains, Jean-François Bernardet for the gift of several *Flavobacterium* sp. and *Chyseobacterium* sp. strains and the CRBIP for also providing us several strains. Laurie Comstock kindly provided the pET16 vector. We thank Rebecca J. Stevick for her support with *in vivo* experiments, Julia Bos for her help with microscopy experiment and, Sebastien Bedu and Nathan Guibert (Institut Pasteur) from the zebrafish facility (Zebrafish projects Hub (Zorgl’hub)) for providing eggs. This work was supported by Institut Pasteur and grants by the French government’s Investissement d’Avenir Program, Laboratoire d’Excellence “Integrative Biology of Emerging Infectious Diseases” (grant n°ANR-10-LABX-62-IBEID). D.P-P. was supported by the Carnot fellowship.

## AUTHOR’S CONTRIBUTIONS

B.A, D.P.-P., S.T.-R., J.B.-B., and F.S. performed the experiments; B.A. D.P.-P. and J.-M.G. designed the experiments and analyzed the data. S.T.-R performed the bioinformatic analysis. J.-M.G. provided resources and funding. B.A. D.P.-P. and J.-M.G. wrote the manuscript.

## REFERENCES

Aoki, S. K., et al. (2010), ‘A widespread family of polymorphic contact-dependent toxin delivery systems in bacteria’, Nature, 468 (7322), 439–42.

Boopathi, S., Liu, D., and Jia, A. Q. (2021), ‘Molecular trafficking between bacteria determines the shape of gut microbial community’, Gut Microbes, 13 (1), 1959841.

Braun, T. F., et al. (2005), ‘Flavobacterium johnsoniae gliding motility genes identified by mariner mutagenesis’, J Bacteriol, 187 (20), 6943–52.

Brennan, A. J., et al. (2010), ‘Perforin deficiency and susceptibility to cancer’, Cell Death Differ, 17 (4), 607–15.

Chatzidaki-Livanis, M., Coyne, M. J., and Comstock, L. E. (2014), ‘An antimicrobial protein of the gut symbiont Bacteroides fragilis with a MACPF domain of host immune proteins’, Mol Microbiol, 94 (6), 1361–74.

Chatzidaki-Livanis, M., Geva-Zatorsky, N., and Comstock, L. E. (2016), ‘Bacteroides fragilis type VI secretion systems use novel effector and immunity proteins to antagonize human gut Bacteroidales species’, Proc Natl Acad Sci U S A, 113 (13), 3627–32.

Chatzidaki-Livanis, M., et al. (2017), ‘Gut Symbiont Bacteroides fragilis Secretes a Eukaryotic-Like Ubiquitin Protein That Mediates Intraspecies Antagonism’, mBio, 8 (6).

Coulthurst, S. (2019), ‘The Type VI secretion system: a versatile bacterial weapon’, Microbiology (Reading), 165 (5), 503–15.

Coyne, M. J. and Comstock, L. E. (2019), ‘Type VI Secretion Systems and the Gut Microbiota’, Microbiol Spectr, 7 (2).

Coyne, M. J., et al. (2019), ‘A family of anti-Bacteroidales peptide toxins wide-spread in the human gut microbiota’, Nat Commun, 10 (1), 3460.

Declercq, A. M., et al. (2013), ‘Columnaris disease in fish: a review with emphasis on bacterium-host interactions’, Vet Res, 44 (1), 27.

Douglas, A. E. (2019), ‘Simple animal models for microbiome research’, Nat Rev Microbiol, 17 (12), 764–75.

Hecht, A. L., et al. (2016), ‘Strain competition restricts colonization of an enteric pathogen and prevents colitis’, EMBO Rep, 17 (9), 1281–91.

Heilbronner, S., et al. (2021), ‘The microbiome-shaping roles of bacteriocins’, Nat Rev Microbiol, 19 (11), 726–39.

Jiang, K., et al. (2024), ‘Bacteroides fragilis ubiquitin homologue drives intraspecies bacterial competition in the gut microbiome’, Nat Microbiol, 9 (1), 70–84.

LaFrentz, B. R., Garcia, J. C., and Shelley, J. P. (2019), ‘Multiplex PCR for genotyping Flavobacterium columnare’, J Fish Dis, 42 (11), 1531–42.

LaFrentz, B. R., et al. (2018), ‘Identification of Four Distinct Phylogenetic Groups in Flavobacterium columnare With Fish Host Associations’, Front Microbiol, 9, 452.

LaFrentz, B. R., et al. (2022), ‘The fish pathogen Flavobacterium columnare represents four distinct species: Flavobacterium columnare, Flavobacterium covae sp. nov., Flavobacterium davisii sp. nov. and Flavobacterium oreochromis sp. nov., and emended description of Flavobacterium columnare’, Syst Appl Microbiol, 45 (2), 126293.

Leshem, A., Liwinski, T., and Elinav, E. (2020), ‘Immune-Microbiota Interplay and Colonization Resistance in Infection’, Mol Cell, 78 (4), 597–613.

Matano, L. M., et al. (2021), ‘Bacteroidetocins Target the Essential Outer Membrane Protein BamA of Bacteroidales Symbionts and Pathogens’, mBio, 12 (5), e0228521.

McEneany, V. L., et al. (2018), ‘Acquisition of MACPF domain-encoding genes is the main contributor to LPS glycan diversity in gut Bacteroides species’, Isme j, 12 (12), 2919–28.

Perez-Pascual, D., et al. (2021), ‘Gnotobiotic rainbow trout (Oncorhynchus mykiss) model reveals endogenous bacteria that protect against Flavobacterium columnare infection’, PLoS Pathog, 17 (1), e1009302.

Peterson, S. B., Bertolli, S. K., and Mougous, J. D. (2020), ‘The Central Role of Interbacterial Antagonism in Bacterial Life’, Curr Biol, 30 (19), R1203–R14.

Pulkkinen, K., et al. (2010), ‘Intensive fish farming and the evolution of pathogen virulence: the case of columnaris disease in Finland’, Proc Biol Sci, 277 (1681), 593–600.

Roelofs, K. G., et al. (2016), ‘Bacteroidales Secreted Antimicrobial Proteins Target Surface Molecules Necessary for Gut Colonization and Mediate Competition In Vivo’, mBio, 7 (4).

Roeselers, G., et al. (2011), ‘Evidence for a core gut microbiota in the zebrafish’, ISME J, 5 (10), 1595–608.

Rosado, C. J., et al. (2008), ‘The MACPF/CDC family of pore-forming toxins’, Cell Microbiol, 10 (9), 1765–74.

Russell, A. B., et al. (2014), ‘A type VI secretion-related pathway in Bacteroidetes mediates interbacterial antagonism’, Cell Host Microbe, 16 (2), 227–36.

Shumaker, A. M., et al. (2019), ‘Identification of a Fifth Antibacterial Toxin Produced by a Single Bacteroides fragilis Strain’, J Bacteriol, 201 (8).

Sorbara, M. T. and Pamer, E. G. (2019), ‘Interbacterial mechanisms of colonization resistance and the strategies pathogens use to overcome them’, Mucosal Immunol, 12 (1), 1–9.

Stagaman, K., Sharpton, T. J., and Guillemin, K. (2020), ‘Zebrafish microbiome studies make waves’, Lab Anim (NY), 49 (7), 201–07.

Stephens, W. Z., et al. (2016), ‘The composition of the zebrafish intestinal microbial community varies across development’, ISME J, 10 (3), 644–54.

Stressmann, F. A., et al. (2021), ‘Mining zebrafish microbiota reveals key community-level resistance against fish pathogen infection’, Isme j, 15 (3), 702–19.

Wang, S., et al. (2024), ‘Microbial collaborations and conflicts: unraveling interactions in the gut ecosystem’, Gut Microbes, 16 (1), 2296603.

Zhu, Y., et al. (2017), ‘Genetic analyses unravel the crucial role of a horizontally acquired alginate lyase for brown algal biomass degradation by Zobellia galactanivorans’, Environ Microbiol, 19 (6), 2164–81.

