## Supplementary Tables and Figures for "A *Chryseobacterium massiliae* pore-forming MACPF domain protein mediates intra and interspecies competition against *Bacteroides*"

**SUPPORTING INFORMATION**

**SUPPLEMENTARY TABLES**

**Table S1 : Strains and plasmids used in this study**

| Strains | Description | Source |
|---|---|---|
| ***Escherichia coli* strains** | | |
| S17 λ pir | Strain used for conjugation | (de Lorenzo and Timmis 1994) |
| BL21 (DE3), pDIA17 | Strain used for expression and purification of a N-terminal His-tagged version of CSAP-1 | |
| ***Chryseobacterium* strains** | | |
| *C. massiliae* I-5479 | Isolated from *Danio rerio* (France) | (Stressmann et al. 2021) |
| *C. massiliae* I-5479 Δ*csap-1* | | This study |
| *C. massiliense* CIP107752T | Isolated from human nose | CRBIP |
| *C. gleum* CIP103039T | Isolated from human vaginal flora (UK) | CRBIP |
| *C. oranimense* CIP109920T | Isolated from raw cow's milk (Israel) | CRBIP |
| *C. indoltheticum* CIP103168T | Isolated from marine mud | CRBIP |
| *C. defluvii* CIP107207T | Isolated from wastewater (Germany) | CRBIP |
| *Chryseobacterium sp.* CIP103055 | Isolated from human sputum (France) | CRBIP |
| *C. gregarium* CIP110171T | Isolated from decaying plant material, litter layer (Germany) | CRBIP |
| *C. piscium* CIP109277T | Isolated from fish (Atlantic ocean) | CRBIP |
| *C. balustinum LMG8329* | Heart blood, dace (*Leuciscus leuciscus*), Dordogne river, Aquitaine, France, 1959 | (Bernardet et al. 2005) |
| *C. scophthalmum LMG13028* | Gills, juvenile turbot (*Scophthalmus maximus*) with hemorrhagic septicemia, Scotland, UK, 1987 | (Bernardet et al. 2005) |
| *C. joostei LMG 18212* | Raw cow milk, Ixopo district, Kwazulu-Natal, South Africa, 1981 | (Bernardet et al. 2005) |
| *Chryseobacterium sp. JIP105/82* | Skin lesion, Koi carp (*Cyprinus carpio*), Ile-de-France, France, 1982 (Japan) | (Bernardet et al. 2005) |
| *Chryseobacterium sp. JIP16/96* | Internal organs, dwarf gourami (*Colisia lalia*), France, 1996 (Singapore) | (Bernardet et al. 2005) |
| *Chryseobacterium sp. JIP17/96* | Kidney, leopard corydoras (*Corydoras julii*), France, 1996 (USA) | (Bernardet et al. 2005) |
| *Chryseobacterium sp. JIP13/00* | Muscle lesion, neon tetra (*Paracheirodon innesi*), France, 2000 (Hong Kong) | (Bernardet et al. 2005) |
| *Chryseobacterium sp. UOF CR2995* | Deep ulcerative dorsal lesion, Atlantic salmon, Finland, 1995 – identified as *C. piscicola* in Ilardi et al, 2009 | (Bernardet et al. 2005) |
| *Chryseobacterium sp. UOF CR4395* | Peduncle lesion, Atlantic salmon, Finland, 1997 - identified as *C. piscicola* in Ilardi et al, 2009 | (Bernardet et al. 2005) |
| *Chryseobacterium sp. JIP 06/01* | Internal organs, rainbow trout, Yvelines, France, 2001 | (Bernardet et al. 2005) |
| *Chryseobacterium sp. SAVU 12/90* | Skin lesion, sea bass (*Dicentrarchus labrax*), Mediterranean coast, France, 1990 | (Bernardet et al. 2005) |

| | | |
|---|---|---|
| *Chryseobacterium sp. LPAA 3120* | Skin lesion, rainbow trout (*Oncorhynchus mykiss*), Brittany, France, 1996 | (Bernardet et al. 2005) |
| *Chryseobacterium sp. LPAA 3230* | Skin lesion, rainbow trout (*Oncorhynchus mykiss*), Brittany, France, 1996 | (Bernardet et al. 2005) |
| *Chryseobacterium sp. LDVH2* | Internal organs, Koi carp, Hérault, France, 2000 (Japan) | (Bernardet et al. 2005) |
| *Chryseobacterium sp. FRGDSA 4580/97* | Siberian sturgeon (*Acipenser baeri*) fry, Landes, France, 1997 – identified as *C. aquaticum* in Loveland-Curtze et al, 2010 | (Bernardet et al. 2005) |
| *Chryseobacterium sp. LDA39 G1966 4a* | Skin lesion, Asian catfish (*Pangasius bocourti*), Kompong Chhnang, Cambodia, 1997 | (Bernardet et al. 2005) |
| *Chryseobacterium sp. FRGDSA 11/99* | Skin ulcer, Koi carp, Landes, France, 1999 | (Bernardet et al. 2005) |
| *Chryseobacterium sp. JG-Tomo* | Liver, rainbow trout, Spain, 2000 | (Bernardet et al. 2005) |
| *Chryseobacterium sp. JG-2* | Skin lesion, rainbow trout, Asturias, Spain, 2000 | (Bernardet et al. 2005) |
| *Chryseobacterium sp. FRGDSA 1502/97* | Kidney, sturgeon (*Acipenser sturio*), Landes, France, 1997 | (Bernardet et al. 2005) |
| *Chryseobacterium sp. LDA39 G1966 13a* | Skin lesion, walking catfish (*Clarias sp.*), Kompong Chhnang, Cambodia, 1997 | (Bernardet et al. 2005) |
| *Chryseobacterium sp. LDVH 1* | Internal organs, goldfish, Hérault, France, 2000 (P. R. of China) | (Bernardet et al. 2005) |
| ***Flavobacterium covae* strains** | | |
| *F. covae* ALG-00-530 | isolated from *Ictalurus punctatus* (USA). | (Olivares-Fuster et al. 2011; Stressmann et al. 2021) |
| *F. covae* JIP 14/00 | isolated from *Paracheirodon innesi* (France). | J.F. Bernardet col. (Stressmann et al. 2021) |
| *F. covae* EK28 | isolated from *Anguilla japonica* (Japan). | J.F. Bernardet col. (Stressmann et al. 2021) |
| *F. covae* AJS1 | isolated from *Poecilia sphenops* (Belgium). | J.F. Bernardet col. (Stressmann et al. 2021) |
| *F. covae* JIP 13/00 | isolated from *Paracheirodon innesi* (Hong Kong). | J.F. Bernardet col. (Stressmann et al. 2021) |
| *F. covae* JIP 02/06(1) | isolated from *Betta splendens* (Singapore). | J.F. Bernardet col. (Stressmann et al. 2021) |
| *F. covae* VB1 | isolated from *Poecilia reticulata* (France). | J.F. Bernardet col. (Stressmann et al. 2021) |
| *F. covae* VB2 | isolated from *Poecilia reticulata* (France). | J.F. Bernardet col. (Stressmann et al. 2021) |
| *F. covae* C#2 | isolated from *Pelteobagrus fulvidraco* (Unknown). | (Li et al. 2017; Stressmann et al. 2021) |
| ***Flavobacterium columnare* strains** | | |
| *F. columnare* FPC666 | isolated from *Misgurnus anguillicaudatus* (Japan). | J.F. Bernardet col. (Stressmann et al. 2021) |

| | | |
|---|---|---|
| *F. columnare* FPC667 | isolated from *Carassius auratus* (Japan). | J.F. Bernardet col. (Stressmann et al. 2021) |
| *F. columnare* LD40 07/2489 | isolated from *Acipenser baeri* (France). | J.F. Bernardet col. (Stressmann et al. 2021) |
| *F. columnare* LVDI 39/I | Unknown | J.F. Bernardet col. (Stressmann et al. 2021) |
| *F. columnare* JIP 17/01 | isolated from *Cyprinus carpio* (France). | J.F. Bernardet col. (Stressmann et al. 2021) |
| *F. columnare* JIP P06/90 | isolated from *Ictalurus melas* (France). | J.F. Bernardet col. (Stressmann et al. 2021) |
| *F. columnare* LVDL 3414/89 | isolated from *Anguilla anguilla* (France). | J.F. Bernardet col. (Stressmann et al. 2021) |
| *F. columnare* UJ H2 | isolated from *Oncorhynchus mykiss* (Finland). | J.F. Bernardet col. (Stressmann et al. 2021) |
| *F. columnare* UJ B259 | isolated from outlet water of a rearing tank with infected *Oncorhynchus mykiss* (Finland). | J.F. Bernardet col. (Stressmann et al. 2021) |
| *F. columnare* UJ FCVK2/872 | isolated from river water above a fish farm (Finland) | J.F. Bernardet col. |
| *F. columnare* Fc4 | isolated from *Oncorhynchus mykiss* (USA). | J.F. Bernardet col. (Stressmann et al. 2021) |
| *F. columnare* Fc7 | isolated from *Oncorhynchus mykiss* (USA). | J.F. Bernardet col. (Stressmann et al. 2021) |
| *F. columnare* JIP P11/91 | isolated from *Oncorhynchus mykiss* (France). | J.F. Bernardet col. (Stressmann et al. 2021) |
| *F. columnare* NCIMB2248 | isolated from *Oncorhynchus tshawytscha* (USA). | J.F. Bernardet col. (Stressmann et al. 2021) |
| *F. columnare* JIP 39/87 | isolated from *Ictalurus punctatus* (France). | J.F. Bernardet col. (Stressmann et al. 2021) |
| *F. columnare* JIP 07/02(1) | isolated from *Cyprinus carpio* (France). | J.F. Bernardet col. (Stressmann et al. 2021) |
| *F. columnare* JIP44/87 (ATCC49512) | isolated from *Samo trutta* (France) | J.F. Bernardet col. (Stressmann et al. 2021) |
| *F. columnare* LVDJ D7461 | isolated from *Oncorhynchus mykiss* (France) | J.F. Bernardet col. |
| *F. columnare* LVDL 2027/89 | isolated from *Acipenser* sp. (France) | J.F. Bernardet col. |
| *F. columnare* IA-S-4 | isolated from walleye (USA) | (Li et al. 2017) |
| *F. columnare* MS-Fc-4 | isolated from *Oncorhynchus mykiss* (USA) | (Bartelme et al. 2018) |
| *F. columnare* LDA39 H4927 | isolated from *Ictalurus melas* (France). | J.F. Bernardet col. (Stressmann et al. 2021) |
| ***Flavobacterium davisii* strains** | | |

| | | |
|---|---|---|
| *F. davisii* SNARCC90 | isolated from *Ictalurus punctatus* (USA). | J.F. Bernardet col. (Stressmann et al. 2021) |
| *F. davisii* CIP109753 | isolated from *Plecoglossus altivelis* (Japan). | CRBIP (Stressmann et al. 2021) |
| *F. ceti CIP109811T* | isolated from whale liver (Spain) | CRBIP |
| *F. branchiophilum* CIP109950 | Isolated from sheatfish (Hungary) | CRBIP |
| *F. psychrophilum* CIP109957 | Isolated from *Oncorhynchus kisutch* (USA) | CRBIP |
| *E. meningoseptica LMG12280* | Isolated from human blood (newborn infant) (USA) | (Bernardet et al. 2005) |
| *B. thetaiotaomicron* VPI5482 | isolated from human feces (CIP104206T) | CRBIP (Xu et al. 2003) |
| *B. ovatus* ATCC8483 | CIP103756T (quality control strain for API products) | CRBIP |
| *B. thetaiotaomicron* CIP103695 | Unknown source (quality control strain for API products) | CRBIP |
| **Plasmids** | | |
| pHimar-Em1 | Plasmid carrying *HimarEm1*, KmR (ErmR) | (Braun et al. 2005) |
| pYT313 | Suicide vector carrying *sacB* under *F. johnsoniae ompA* promoter, AmpR (ErmR). | (Zhu et al. 2017) |
| pET16 | Vector used to construct a N-terminal His-tagged version of CSAP-1 | Gift from Comstock lab |

**Table S2 : Primers used in this study**

| | Name | Sequence (5'->3') |
|---|---|---|
| pYT313 [(1)] vector amplification | pYT313-1 | TTCCTCGCTCACTGACTCGC |
| | pYT313-2 | TGACCATGATTACGCCAAGC |
| *csap-1* deletion mutant in *C. massiliae* | *csap* -UP-For | GCTTGGCGTAATCATGGTCAGCACAATATACCATTCCCCC |
| | *csap* -UP-Rev | ACTTTTTTACAGGGCATACCTTAAAGTTTTTCTCATAAA |
| | *csap* -DOWN-For | TATGCCCTGTAAAAAAGT |
| | *csap* -DOWN-Rev | GTCAGTGAGCGAGGAACGTTGACATCTTTTCCCG |
| to confirm *csap-1* deletion in *C. massiliae* | *csap* - ext5 | CGGAAGCAGGAGCAGAAAAGG |
| | *csap* - ext3 | CACTCCCAAAATAACGGTCGC |
| Construction of His-tagged CSAP-1 | pET16- *csap* -For | GCTTTTTATGCCCTGTAACTAGCATAACCCCTTGGG |
| | pET16- *csap* -Rev | CCAAAATTTCATTTGTACTACGACCTTCGATATGGCCG |
| | *csap* - pET16-For | CGGCCATATCGAAGGTCGTAGTACAAATGAAATTTTGG |
| | *csap* - pET16-Rev | CCCAAGGGGTTATGCTAGTTACAGGGCATAAAAAGC |
| Multiplex PCR for genotyping *F. columnare* [(2)] | GG-fwd | ACRGGRGATAAAGCAGAASA |
| | GG1-rev | GACTTTTGTGTTGAAACGG |
| | GG2-rev | AAGAAAATAGGGGAGAGG |
| | GG3-rev | CAAGTTTCGTTATGATGAGG |
| | GG4-rev | TCCAAAAAGTCCGMAATC |

(1) (Zhu et al. 2017)
(2) (LaFrentz et al. 2019)

**Table S3**: Genotyping 33 isolates using the multiplex PCR to revise the species name[1].

| Name | Genetic group | Species name | Reference |
|---|---|---|---|
| ALG-00-530 | 2 | *F. covae* | (LaFrentz et al. 2022) |
| JIP 14/00 | 2 | *F. covae* | This study |
| EK28 | 2 | *F. covae* | This study |
| AJS1 | 2 | *F. covae* | This study |
| JIP 13/00 | 2 | *F. covae* | This study |
| JIP 02/06(1) | 2 | *F. covae* | This study |
| VB1 | 2 | *F. covae* | This study |
| VB2 | 2 | *F. covae* | This study |
| C#2 | 2 | *F. covae* | (LaFrentz et al. 2022) |
| FPC666 | 1 | *F. columnare* | This study |
| FPC667 | 1 | *F. columnare* | This study |
| LD40 07/2489 | 1 | *F. columnare* | This study |
| LVDI 39/I | 1 | *F. columnare* | This study |
| JIP 17/01 | 1 | *F. columnare* | This study |
| JIP P06/90 | 1 | *F. columnare* | This study |
| LVDL 3414/89 | 1 | *F. columnare* | This study |
| UJ H2 | 1 | *F. columnare* | This study |
| UJ B259 | 1 | *F. columnare* | This study |
| UJ FCVK2/872 | 1 | *F. columnare* | This study |
| Fc4 | 1 | *F. columnare* | This study |
| Fc7 | 1 | *F. columnare* | This study |
| JIP P11/91 | 1 | *F. columnare* | This study |
| NCIMB2248 | 1 | *F. columnare* | This study |
| JIP 39/87 | 1 | *F. columnare* | This study |
| JIP 07/02(1) | 1 | *F. columnare* | This study |
| JIP44/87 (ATCC49512) | 1 | *F. columnare* | (LaFrentz et al. 2018) |
| LVDJ D7461 | 1 | *F. columnare* | This study |
| LVDL 2027/89 | 1 | *F. columnare* | This study |
| IA-S-4 | 1 | *F. columnare* | (LaFrentz et al. 2022) |
| MS-Fc-4 | 1 | *F. columnare* | This study |
| LDA39 H4927 | 1 | *F. columnare* | This study |
| SNARCC90 | 3 | *F. davisii* | This study |
| CIP109753 | 3 | *F. davisii* | This study |

[1] (LaFrentz et al. 2018; LaFrentz et al. 2019; LaFrentz et al. 2022)

**SUPPLEMENTARY FIGURES**

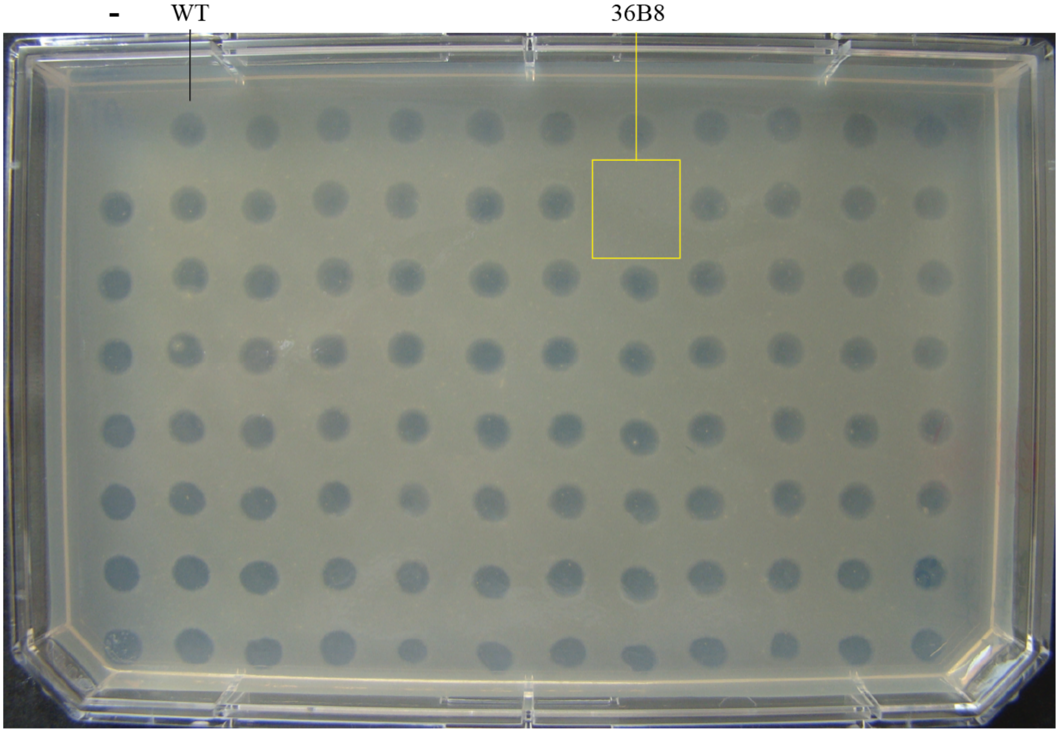


**Figure S1: Identification of *C. massiliae* transposon mutant unable to inhibit *F. covae*$^{ALG}$ growth.**

A plate representative of the screening performed to identify transposon mutants whose supernatant does not display any growth inhibitory activity against *F. covae*$^{ALG}$. *F. covae*$^{ALG}$ was grown as an overlay and 2 µL of filtered supernatant from each transposon mutant was spotted on top of it. Negative control (-) corresponded to 2 µL of TYES medium; WT: 2 µL of supernatant from WT strain of *C. massiliae* (positive control). The yellow square indicates one (36B8) of the 4 identified transposon mutants unable to inhibit the growth of *F. covae*$^{ALG}$.

**A**

| | BSAP-1 | BSAP-2 | BSAP-3 | BSAP-4 | CSAP-1 |
|---|---|---|---|---|---|
| BSAP-1 | 100 | 27,05 | 34,89 | 24,07 | 29,63 |
| BSAP-2 | 27,05 | 100 | 32,11 | 23,02 | 23,11 |
| BSAP-3 | 34,89 | 32,11 | 100 | 22,17 | 26,62 |
| BSAP-4 | 24,07 | 23,02 | 22,17 | 100 | 18,46 |
| CSAP-1 | 29,63 | 23,11 | 26,62 | 18,46 | 100 |

**B**

```
BSAP-4      NDYGTHVLVDFVLGGKLSVVTSAIDHSDNKDELFK----FS--TKIFKIISASASSSISK      281
CSAP-1      QYYGTHILTDIYTGGKLDVMFRSETTNENRKN----ASEVGVKVSALNIFNINVNNNVDT      244
BSAP-1      QHYGTHVLTDITLGGRITVLYRSSINTSKKTA----TVEAGCASGIKNMFNLSVDGHYDQ      238
BSAP-2      EEFGTHILIDFNVGGRLSIFYKSTITDNLKVESKTKIAKGG-ITGAIKTVNLSFSGSSTT      267
BSAP-3      AKYGTHVLTNITVGGSYIAYYKSAIIEENTGTEKKQTVSAG-AKYNMSKVGLDFDGTWST      253
              :***:* ::  **       :    .            .      . .. . ..

BSAP-4      S----NY--LKNTSLTILQAGGSEIQAVKRYIEEDGTINSDIFNYKEWIKSINKETSVLV      335
CSAP-1      SS----ASKNFNRTLSYRSYGGDPTKSLLGTISMDQS-TP-SVNIANWQSSCTPENSVFV      298
BSAP-1      TLVKDNSE----QEIVYRTEGGDPSRALIGQLNYDSK-NPSVIDISSWQQSCDDNNMTLV      293
BSAP-2      TEVEQYQRRNSNWSCNVNMYGGQHDGHTI-TITSDGA-TNHTFNLSGWQQSVDKTHCVLT      325
BSAP-3      TTITEANKKNSDWTCTIQCVGGTTSGTTI-TLAPNQG-PTTTINIGAWSQSVDDTHSRLV      311
            :                   **         :  :       .:   * .*       :.
```

**Figure S2: Sequence comparison of CSAP-1 with *Bacteroides* BSAP proteins displaying intraspecies inhibitory activity.**

**(A)** Percent identity matrix created by Clustal2.1 on Uniprot website. **(B)** Alignment of the conserved residues of MACPF domains of BSAP-1 to -4 and CSAP-1. The conserved MACPF motif (Y/W-G-T/S-H-F/Y-$X_6$-GG) and the residue W (boxed in red) are critical residues mapped in relation to other MACPF containing proteins (Chatzidaki-Livanis et al. 2014).

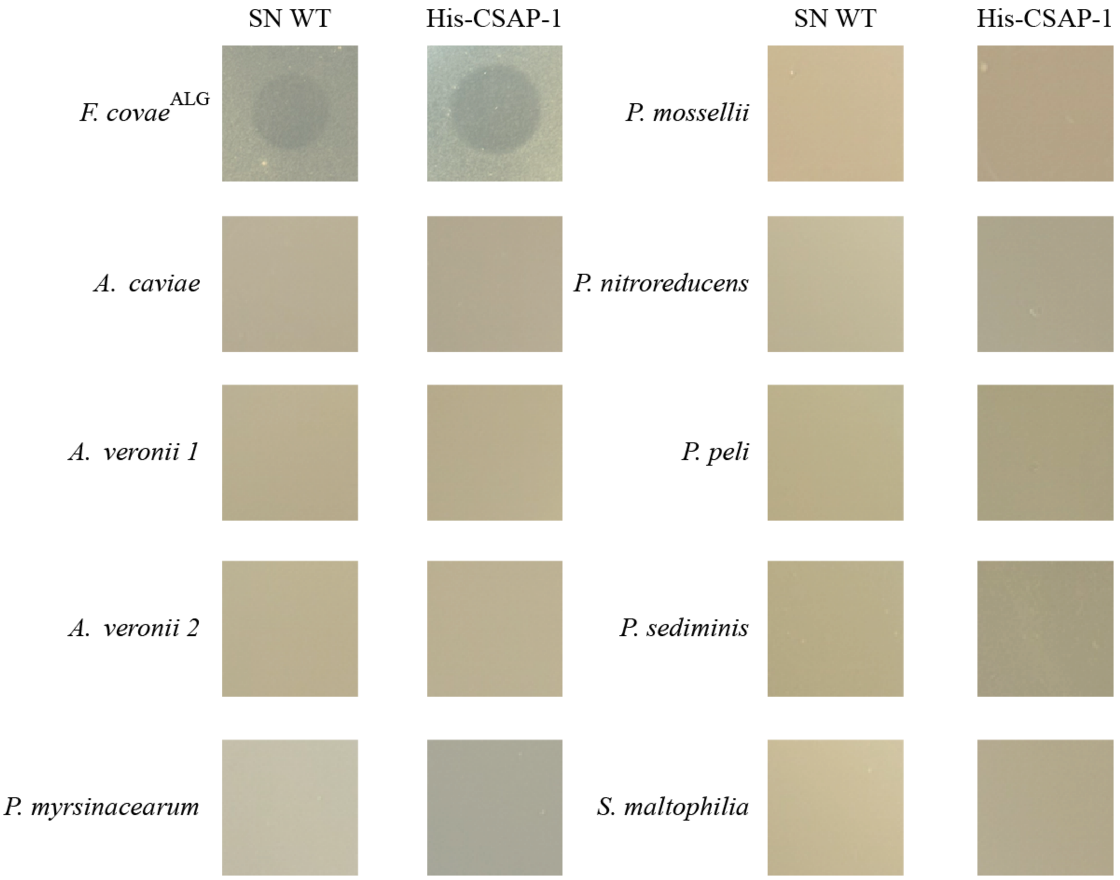


**Figure S3: CSAP-1 is inactive against nine proteobacteria from the zebrafish larvae microbiota.**

Overlay assay showing that CSAP-1 from *C. massiliae* does not inhibit the growth of the nine proteobacteria isolated from the zebrafish larvae microbiota (Stressmann et al. 2021). 10 µl of filtered supernatant of a *C. massiliae* culture or purified His-CSAP-1 were dropped on overlaid bacteria.

A

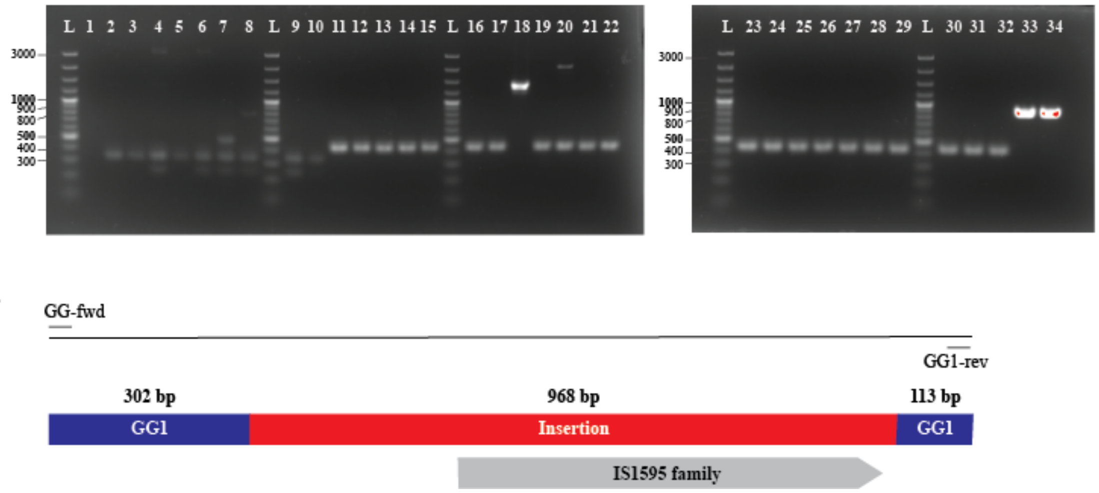


B

**Figure S4: Multiplex PCR for genotyping *F. columnare* and revision of species designation** (A)Gel electrophoresis of PCR products amplified from gDNA using the multiplex PCR conditions described in (LaFrentz et al. 2019). Lane L: GeneRuler 100 bp Plus DNA ladder (Thermo Scientific, SM0321); Lane 1: negative control (no template); Lane 2: ALG-00-530; Lane 3: JIP14/00; Lane 4: EK28; Lane 5: AJS1; Lane 6: JIP 13/00; Lane 7: JIP 02/06(1); Lane 8: VB1; Lane 9: VB2; Lane 10: C#2; Lane 11: FPC666; Lane 12: FPC667; Lane 13: LD40 07/2489; Lane 14: LVDI 39/I; Lane 15: JIP 17/01; Lane 16: JIP P06/90; Lane 17: LVDL 3414/89; Lane 18: UJ H2; Lane 19: UJ B259; Lane 20: UJ FCVK2/872; Lane 21: Fc4; Lane 22: Fc7; Lane 23: JIP 11/91; Lane 24: NCIMB2248; Lane 25: JIP 39/87; Lane 26: JIP 07/02(1); Lane 27: JIP44/87; Lane 28: LVDJ D7461; Lane 29: LVDL 2027/89; Lane 30: IA-S-4; Lane 31: MS-Fc-4; Lane 32: LDA39 H4927; Lane 33: SNARCC90; Lane 34: CIP109753. Sizes corresponding to each species were 415, 320 and 894 bp for respectively *F. columnare* (genetic group 1), *F. covae* (genetic group 2) and *F. davisii* (genetic group 3). No *F. oreochromis* (genetic group 4) were identified among our 33 isolates used in our study. (B)Multiplex PCR target sequence in *F. columnare* UJH2. The amplicon size for the strain UJH2 being higher than expected (1.3kb), the PCR product was sequenced and compared to the expected sequences depending on each genetic group. The alignment revealed that the sequence corresponds to genetic group 1 with an insertion (IS1595 family).

**Supplementary references**

Bartelme, R. P., et al. (2018), 'Draft Genome Sequence of the Fish Pathogen Flavobacterium columnare Strain MS-FC-4', *Genome Announc,* 6 (20).

Bernardet, J. F., et al. (2005), 'Polyphasic study of Chryseobacterium strains isolated from diseased aquatic animals', *Syst Appl Microbiol,* 28 (7), 640-60.

Braun, T. F., et al. (2005), 'Flavobacterium johnsoniae gliding motility genes identified by mariner mutagenesis', *J Bacteriol,* 187 (20), 6943-52.

Chatzidaki-Livanis, M., Coyne, M. J., and Comstock, L. E. (2014), 'An antimicrobial protein of the gut symbiont Bacteroides fragilis with a MACPF domain of host immune proteins', *Mol Microbiol,* 94 (6), 1361-74.

de Lorenzo, V. and Timmis, K. N. (1994), 'Analysis and construction of stable phenotypes in gram-negative bacteria with Tn5- and Tn10-derived minitransposons', *Methods Enzymol,* 235, 386-405.

LaFrentz, B. R., Garcia, J. C., and Shelley, J. P. (2019), 'Multiplex PCR for genotyping Flavobacterium columnare', *J Fish Dis,* 42 (11), 1531-42.

LaFrentz, B. R., et al. (2018), 'Identification of Four Distinct Phylogenetic Groups in Flavobacterium columnare With Fish Host Associations', *Front Microbiol,* 9, 452.

LaFrentz, B. R., et al. (2022), 'The fish pathogen Flavobacterium columnare represents four distinct species: Flavobacterium columnare, Flavobacterium covae sp. nov., Flavobacterium davisii sp. nov. and Flavobacterium oreochromis sp. nov., and emended description of Flavobacterium columnare', *Syst Appl Microbiol,* 45 (2), 126293.

Li, N., et al. (2017), 'The Type IX Secretion System Is Required for Virulence of the Fish Pathogen Flavobacterium columnare', *Appl Environ Microbiol,* 83 (23).

Olivares-Fuster, O., et al. (2011), 'Adhesion dynamics of Flavobacterium columnare to channel catfish Ictalurus punctatus and zebrafish Danio rerio after immersion challenge', *Dis Aquat Organ,* 96 (3), 221-7.

Stressmann, F. A., et al. (2021), 'Mining zebrafish microbiota reveals key community-level resistance against fish pathogen infection', *Isme j,* 15 (3), 702-19.

Xu, J., et al. (2003), 'A genomic view of the human-Bacteroides thetaiotaomicron symbiosis', *Science,* 299 (5615), 2074-6.

Zhu, Y., et al. (2017), 'Genetic analyses unravel the crucial role of a horizontally acquired alginate lyase for brown algal biomass degradation by Zobellia galactanivorans', *Environ Microbiol,* 19 (6), 2164-81.
